# A RHNO1-ATR/Chk1 positive feedback loop sustains the DNA replication stress response

**DOI:** 10.64898/2026.05.22.727300

**Authors:** Niphat Jirapongwattana, Carley M. Conover, Catalina Trujillo Jaramillo, Kishore K. Chiruvella, Joonyoung Her, Gargi Ghosal, Samuel F. Bunting, Dale A. Ramsden, Adam R. Karpf

## Abstract

Elevated DNA replication stress is a common feature of cancer cells, rendering them dependent on ATR/Chk1 signaling, which controls the DNA replication stress checkpoint, for survival. Although the activation of ATR/Chk1 signaling is well-established, how this signal is maintained until replication stress is fully resolved is less understood. Here, we investigated the roles of RAD9–HUS1–RAD1 interacting nuclear orphan 1 (RHNO1) in cancer progression and its involvement in maintaining ATR/Chk1 signaling during the DNA replication stress response. We demonstrate that depletion of RHNO1 significantly inhibits cancer cell proliferation *in vitro* and tumor growth *in vivo*. Mechanistically, we show that, while RHNO1 is dispensable for initial ATR/Chk1 signal activation, it is upregulated and stabilized following replication stress. RHNO1 is required to sustain ATR/Chk1 signaling, prevent premature checkpoint collapse, and suppress genomic instability. Under basal conditions, RHNO1 is rapidly degraded by the proteasome, however, RHNO1 is phosphorylated and stabilized following DNA replication stress. Stabilization of RHNO1 is mediated by ATR/Chk1 signaling which promotes RHNO1 localization to stressed replication forks marked by phosphorylated RPA32. Together, our data reveals a novel positive feedback loop wherein ATR/Chk1 signaling activation stabilizes RHNO1, which, in turn, is required to sustain the signal and thus the replication stress response. This work identifies RHNO1 as a key component in the cellular replication stress response and highlights its potential as a therapeutic target for tumor cells reliant on ATR/Chk1 signaling.

## Introduction

Cancer cells are characterized by dysregulated cell proliferation and DNA replication, commonly leading to the slowing or stalling of replication forks, known as DNA replication stress^1,2^. Unresolved replication stress can lead to replication fork collapse and DNA double-strand breaks, an extremely toxic form of genomic instability^3,4^. Therefore, it is imperative for cancer cells to properly mitigate replication stress to ensure their survival.

Ataxia Telangiectasia and Rad3-related (ATR) kinase is the master regulator of the replication stress response^5^. Activation of ATR and its key downstream kinase checkpoint kinase 1 (Chk1) promotes cell cycle arrest, inhibition of new origin firing, stabilization of replication forks, and the DNA damage repair (DDR) response, collectively known as the DNA checkpoint^5–8^. ATR/Chk1-mediated replication stress responses by slow down DNA replication, allowing cells to resolve replication stress before fork collapse and DNA damage ensues. Importantly, cancer cells have elevated replication stress, which results from oncogene expression as well as from extrinsic factors including oxidative stress and chemotherapy exposure, creating a strong reliance on ATR/Chk1 signaling for cancer cell survival^9–11^. While the canonical mechanism of ATR/Chk1 activation is well established^12–14^, how this signaling is sustained chronically until replication stress is fully resolved is less understood.

RAD9–HUS1–RAD1 interacting nuclear orphan 1 (RHNO1, previously called C12orf32 and RHINO) was first identified as a component of DNA damage response machinery, binding to the 9-1-1 and TopBP1 complex on RPA32-coated single-stranded DNA (ssDNA) to facilitate ATR activation^15^. Subsequent studies revealed that RHNO1 forms a stoichiometric heterotetrameric complex with 9-1-1 during ATR/Chk1 activation^16,17^. Crucially, previous studies have shown that the 9-1-1 complex can be loaded onto chromatin and interact with TopBP1 even in the absence of RHNO1^16^. However, in this condition, the ATR/Chk1 signal is partially abrogated ^15,16,18^. These findings raise a question regarding the exact function of RHNO1 in ATR/Chk1 signaling as the 9-1-1/TopBP1 complex alone is sufficient for ATR activation.

In this study, we investigated the function of RHNO1 during the DNA replication stress response and how it affects cancer cell and tumor growth and survival. We observed that RHNO1-depletion impairs ovarian cancer growth and survival both *in vitro* and *in vivo*. Mechanistically, we show that RHNO1 functions not in ATR/Chk1 checkpoint activation but rather in its maintenance. RHNO1 is phosphorylated by ATR/Chk1 signaling in cells undergoing DNA replication stress, which subsequently protects RHNO1 from proteasomal degradation. Post-translational stabilization of RHNO1 leads to increased RHNO1 protein levels and its localization at stalled replication forks. Based on our findings, we propose the existence of a positive feedback loop between RHNO1 and ATR/Chk1 signaling where RHNO1 is dispensable for the initiation of ATR/Chk1 signaling but is critical for sustaining the checkpoint signal. From a therapeutic perspective, we hypothesize that disrupting this feedback loop by targeting RHNO1 pharmacologically could lead to premature checkpoint collapse and increased genomic instability in cells experiencing ongoing replication stress.

## Results

### RHNO1 depletion reduces oncogenic phenotypes, impairs the replication stress response, and increases DNA damage in ovarian cancer cells

To investigate the role of RHNO1 in ovarian cancer cell proliferation, we generated constitutive RHNO1 knockdown OVCAR8 cell lines (OVCAR8 RH130 and RH133). RHNO1 depletion significantly reduced cell proliferation in comparison to the non-silencing control (OVCAR8 NS) (**Fig. 1A**). Consistent with the cell proliferation assay, both colony-formation and soft-agar colony-formation (2D and 3D survival) assays demonstrated a significantly reduced number of colonies with RHNO1 knockdown (**Fig. 1B-D**). Given the known function of RHNO1 in the ATR/Chk1 signaling pathway, we next assessed RHNO1 mRNA levels under replication stress conditions using hydroxyurea (HU) treatment, which inhibits ribonucleotide reductase, depleting nucleotide pools^19^. After 24 hours of 5 mM HU treatment, the *RHNO1* mRNA level in the non-silencing control (NS) was significantly increased compared to the untreated counterpart, while in RHNO1 knockdown lines, *RHNO1* remained suppressed (**Fig. 1E**).

**Figure 1.**
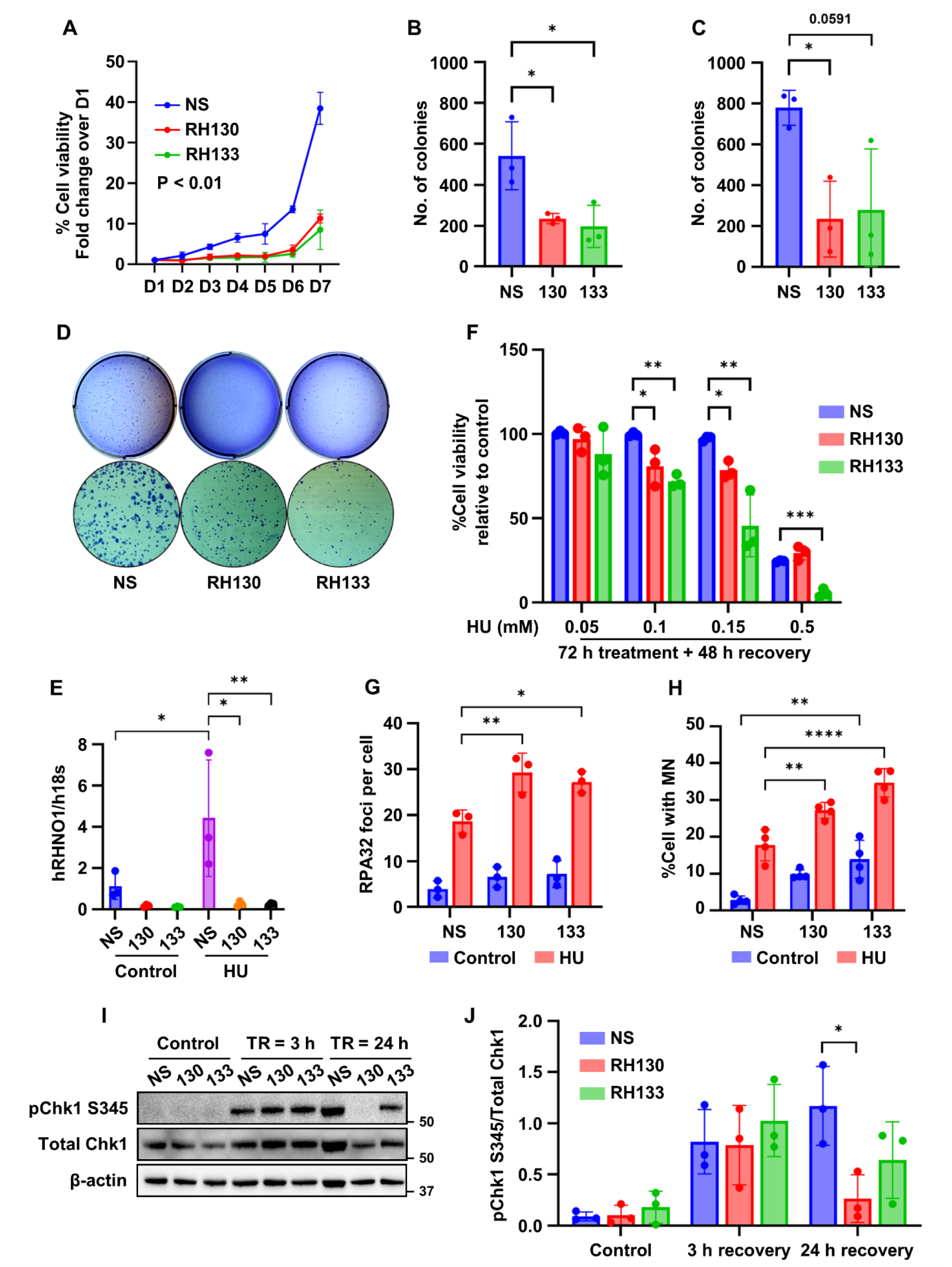
RHNO1 depletion inhibits ovarian cancer cell proliferation and impairs the replication stress response. **A** Cell proliferation rates of OVCAR8 cells stably express non-silencing (NS) or RHNO1-specific shRNA (RH130 and RH133) monitored over 7 days. **B** and **C** Quantification of colonies in colony-formation assay (**B**) and soft-agar colony-formation assay (**C**). **D** Representative images of crystal violet stained colonies from **B** (top panel) and **C** (bottom panel). **E** RHNO1 mRNA levels in OVCAR8 NS and RHNO1 knockdown cells following 24 hours treatment with 5 mM hydroxyurea (HU). **F** Cell viability of OVCAR8 cells relative to control following various concentration of HU treatment for 72 hours followed by 48 hours recovery. **G** Quantification of RPA32 foci per cell after 5 hours treatment of 2.5 mM HU followed by 24 hours recovery. **H** Percentage of cells with micronuclei after 5 hours of 5 mM HU treatment and 48 hours recovery. **I** Western blotting of ATR/Chk1 signaling pathway after 3 hours of 5 mM HU treatment followed by 3 and 24 hours (time to recovery TR) after washout. **J** Quantification of pChk1 S345 normalized by total Chk1 from figure **I**. Data is presented as mean ± SD of at least three independent experiments. Statistical significance was determined using one-way ANOVA with Tukey post-hoc test (*p < 0.05, **p < 0.01, ***p < 0.001, ****p < 0.0001).

Next, we investigated the impact of RHNO1 depletion on the sustained replication stress response; cells were incubated with various concentrations of HU for 72 hours and allowed to recover for 48 hours before evaluating cell viability. Both RHNO1 knockdown lines significantly increased OVCAR8 cell sensitivity to 0.1 and 0.15 mM HU treatment in comparison to the NS counterpart (**Fig. 1F**). The slight variance in viability at the highest 0.5 mM HU treatment likely reflects minor differences in residual RHNO1 protein levels or general stress tolerance between the two knockdown lines. Consistent with these findings, there were significant elevations of RPA32 foci, a marker of ongoing replication stress, in RHNO1 knockdown OVCAR8 cell lines treated with 2.5 mM HU for 5 hours and allowed to recover for 24 hours (**Fig. 1G, Supplementary Fig. 1A**). Interestingly, despite the significantly increased RPA32 foci, the phosphorylated RPA32 serine 33 (pRPA32 S33), a downstream signaling marker of ATR, was lower in RHNO1 knockdown cells compared to NS control (**Supplementary Fig. 1B**)^13^. The increase of RPA32 but not pRPA32 S33 foci in RHNO1 knockdown cells suggests an impaired ATR signal under replication stress conditions, as S33 is a target of ATR phosphorylation^13^.

A defective replication stress response can result in micronuclei (MN) formation, resulting from the unrepaired single-stranded DNA breaks progressing into the late cell cycle phases ^20^. Notably, OVCAR8 cells sustaining RHNO1 knockdown, even without HU treatment, showed a significant increase of MN-containing cells, and MN formation in these cells was further exacerbated by 5 hours of 5 mM HU treatment and 48 hours recovery (**Fig. 1H, Supplementary Fig. 2A**).

Intriguingly, OVCAR8 control and RHNO1 knockdown cells demonstrated comparable ATR/Chk1 signaling activation after 3 hours recovery from 5 mM HU treatment, regardless of RHNO1 levels (**Fig. 1I** and **1J**). However, when cells were allowed to recover for 24 hours after HU washout, the NS control cells still retained elevated phosphorylated Chk1 serine 345 (pChk1 S345), while the RHNO1-depleted cells showed premature loss of pChk1 S345 (**Fig. 1I** and **J**). The differential decreased pChk1 S345 levels observed between RH130 and RH133, may be due to the clonal variation or slight differences in residual RHNO1 protein levels, which we are not able to assess in this system due to lack of a validated RHNO1 primary antibody. These findings suggest that RHNO1 is not required for the initial activation of ATR/Chk1 signaling but instead contributes to sustaining ATR/Chk1 signaling during ongoing replication stress.

### RHNO1 depletion impairs polymerase theta-mediated end joining (TMEJ) in ovarian cancer cells

It was recently reported that, in addition to its role in the replication stress response, RHNO1 promotes polymerase theta-mediated end joining (TMEJ) during M phase^21^. Therefore, to further validate our OVCAR8 RHNO1 knockdown model system, we determined whether RHNO1 knockdown impairs TMEJ in these cells. Notably, RHNO1 depletion significantly reduced the microhomology deletion (MHD) and templated insertions (TINs) caused by TMEJ pathway (**Supplementary Fig. 3A**-**C**). Interestingly, addition of ART558, a DNA polymerase θ inhibitor, can further reduce the TMEJ activity in RHNO1-depleted cells, suggesting a residue TMEJ function despite the absence of RHNO1 (**Supplementary Fig. 3D**). While these data support that defective TMEJ may contribute to the baseline genomic instability seen in RHNO1 knockdown cells, the premature collapse of ATR/Chk1 signaling following HU treatment was the focus of the current investigation.

### RHNO1 depletion inhibits ovarian cancer xenograft growth and prolongs mouse survival

To evaluate the effects of RHNO1 depletion in ovarian cancer in vivo, we tagged RHNO1 knockdown OVCAR8 cells with luciferase and intraperitoneally transplanted these cells into immunocompromised mice. RHNO1-depleted tumors exhibited significantly reduced tumor growth (**Fig. 2A**). Notably, the reduced tumor growth was accompanied by lower frequencies of mice with ascites, a marker of ovarian cancer progression, at the time of euthanasia (**Fig. 2B**). Furthermore, the mice transplanted with RHNO1-depleted cells demonstrated significantly prolonged overall survival time compared to the NS control group (**Fig. 2C**). Together, these data are consistent with an important role for RHNO1 in promoting ovarian cancer cell growth in vivo (**Fig. 2C**).

**Figure 2.**
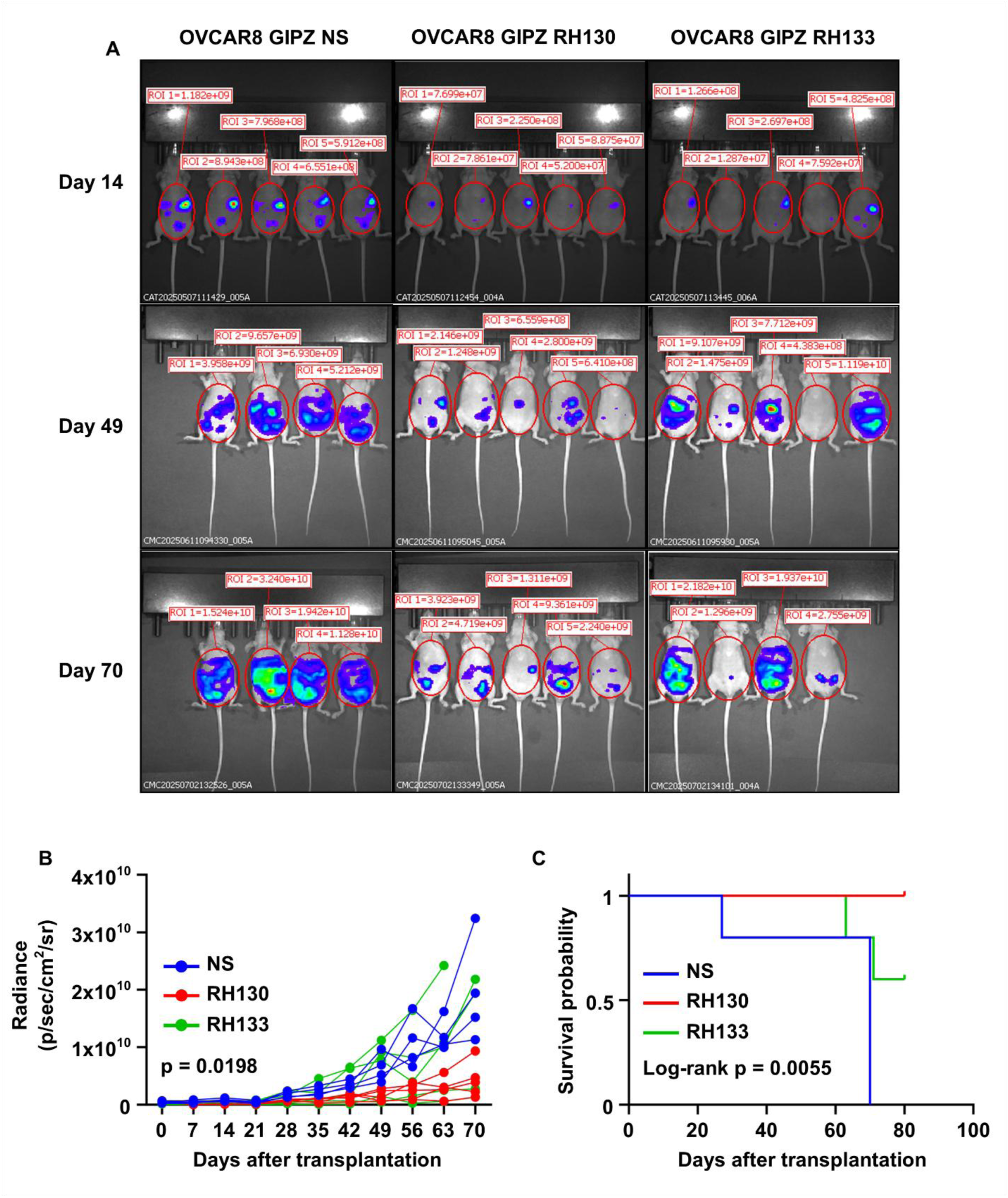
RHNO1 depletion inhibits tumor growth and prolongs survival in an ovarian cancer xenograft model**. A** Representative bioluminescence images of nude mice intraperitoneally transplanted with luciferase-expressing OVCAR8 NS, RH130 or RH133 cells at days 14, 49, and 70 post-transplantations. **B** Quantification of tumor growth over 70 days post-transplantation. **C** Kaplan-Meier survival analysis comparing mice transplanted with OVCAR8 NS, RH130, and RH133. Statistical significance was determined using two-way ANOVA (**B**), and log-rank test (**C**).

### Replication stress-induced phosphorylation inhibits RHNO1 proteolytic degradation

As mentioned, studying RHNO1 is challenging due to the lack of available primary antibodies capable of recognizing endogenous RHNO1 (data not shown). Thus, to allow investigation of the functions of endogenous RHNO1, we generated C-terminal HiBiT-tagged RHNO1 HEK293T cells using CRISPR-Cas9 knock-in (**Supplementary Fig. 4A** and **B**). HiBiT tagging is advantageous because proteins can be detected either by antibody-based methods or luminescence^22^. We first confirmed that the RHNO1-HiBiT protein can be specifically detected using the HiBiT luminescence assay and showed that luminescence correlated with cell number (**Supplementary Fig. 4C** and **D**). Next, we tracked the kinetics of RHNO1 protein expression under replication stress induced by 2.5 mM HU and found that RHNO1 protein gradually increased over time, achieving statistically significant levels only after 6 hours of treatment (**Fig. 3A**). To investigate a possible correlation between RHNO1 levels and a DNA replication stress response, we next collected cell lysates at 2, 6, and 24 hours after either 2.5 mM HU or 5 µM etoposide treatment and conducted western blotting. We found that RHNO1 only reached detectable levels at 6 hours after treatment, consistent with the luminescence assay, and was further increased at 24 hours after treatment (**Fig. 3B**). Strikingly, in addition to its gradual accumulation, there was an increase in molecular weight of RHNO1 under replication stress conditions (**Fig. 3B**). Comparing the kinetics of RHNO1 induction to established replication stress response markers revealed a distinct temporal pattern: while the acute phase marker pChk1 S345 was robustly activated as early as 2 hours and persisted throughout the treatment, RHNO1 accumulation was delayed. Instead, the kinetics of RHNO1 accumulation and modification closely mirrored that of pRPA32 S33 (**Fig. 3B**). The induction of pChk1 S345 at the earlier timepoints supports our findings in OVCAR8 RHNO1 knockdown cells that ATR/Chk1 signaling can be activated in the absence of RHNO1. More importantly, the data suggests that RHNO1 may be post-translationally modified by activated ATR/Chk1 signaling, which potentially plays a role in sustaining the response.

**Figure 3.**
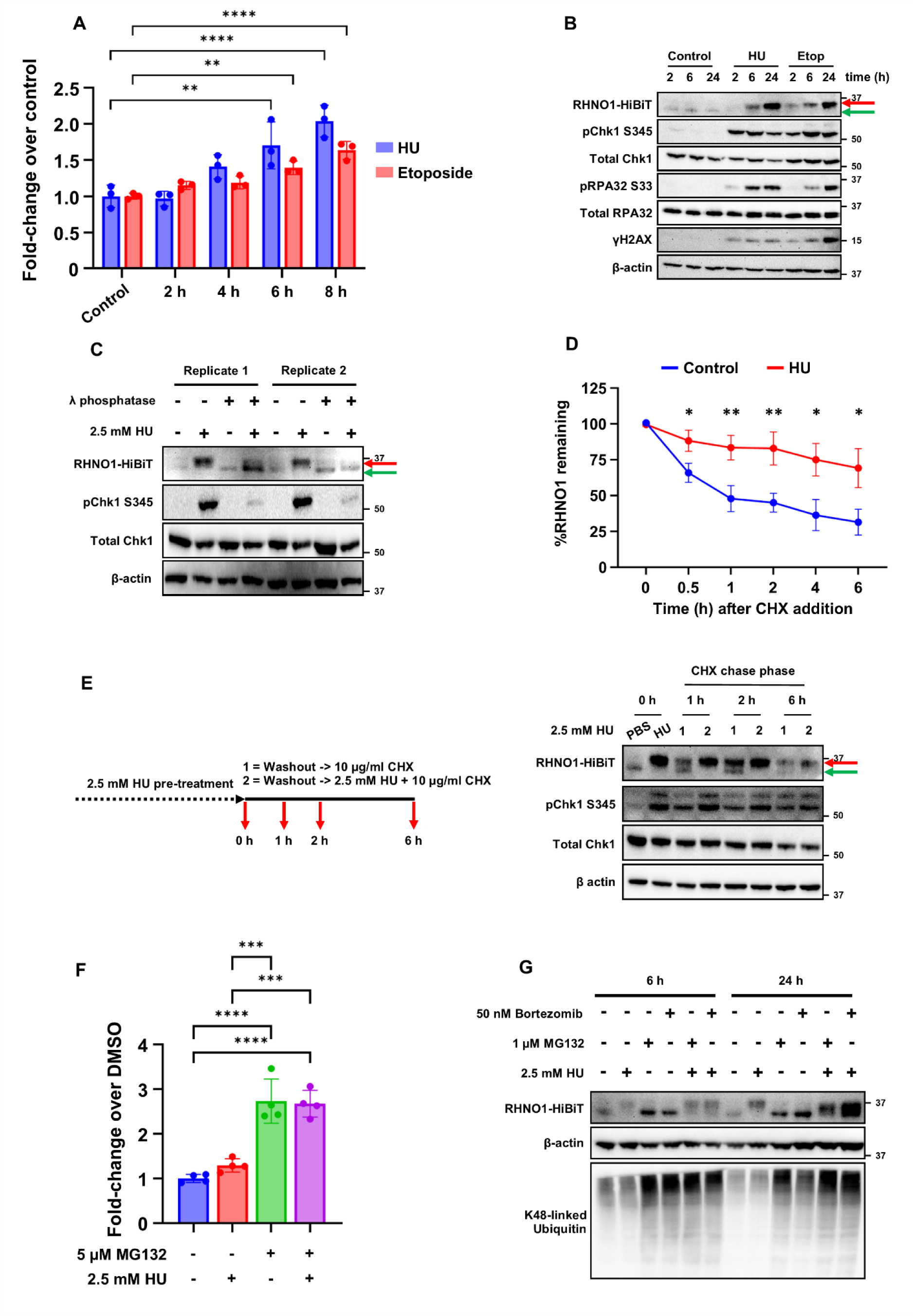
RHNO1 is phosphorylated and stabilized in replication stress conditions**. A** The kinetics of RHNO1 protein measured by HiBiT luminescence in HEK293T RHNO1-HiBiT cells treated with 2.5 mM HU or 5 µM etoposide over 8 hours. **B** Western blotting of RHNO1-HiBiT on a time-course incubation of 2.5 mM HU or 5 µM etoposide. Phosphorylated and unphosphorylated RHNO1 are noted by red and green arrows, respectively. **C** Western blot analysis of 24 hours 2.5 mM HU-treated HEK293T RHNO1-HiBiT cell lysates incubated with or without 400 U of λ phosphatase. **D** Cycloheximide (CHX) chase assay of RHNO1-HiBiT in control and 18 hours 2.5 mM HU-pretreated cells over 6 hours analyzed by HiBiT luminescence assay. **E** Western blot analysis of RHNO1-HiBiT phosphorylation and stability during a CHX chase. Following a 24 hours pre-treatment with 2.5 mM HU, cells were chased with CHX under either HU washout conditions (1) or continuous HU treatment (2). **F** Quantification of RHNO1-HiBiT protein following 5 µM MG132 with or without 2.5 mM HU treatment for 6 hours by HiBiT luminescence assay. **G** Western blot analysis of RHNO1-HiBiT after 6 and 24 hours treatment of HU with or without MG132 or Bortezomib. Data is presented as mean ± SD of at least three independent experiments. Statistical significance was determined using unpaired Student’s t test two-tailed (**D**), and one-way ANOVA with Tukey post-hoc test (**A** and **F**) (*p < 0.05, **p < 0.01, ***p < 0.001, ****p < 0.0001).

Based on the molecular weight change of RHNO1 under replication stress, we hypothesized that this was a phosphorylation modification. To test this, we incubated HU-treated cell lysates with λ phosphatase prior to RHNO1 detection by western blotting. Notably, under these conditions, the high molecular weight form of RHNO1 shifted to the lower molecular weight form, strongly suggesting that RHNO1 is phosphorylated under replication stress conditions (**Fig. 3C**). To determine whether RHNO1 accumulation is specific to HU and etoposide treatment, or is more generalizable, we treated HEK293T RHNO1-HiBiT cells with a panel of DNA-damaging agents. In line with the previous findings, agents that induce high levels of pChk1 S345 and pRPA32 S33 generally promote either RHNO1 accumulation and/or phosphorylation (**Supplementary Fig. 5A**).

Phosphorylation is a common mechanism of cells to regulate protein levels, by influencing protein stability and degradation^23,24^. Therefore, we hypothesized that the phosphorylation of RHNO1 seen under replication stress may result in RHNO1 stabilization and/or accumulation. To test this hypothesis, first we performed a cycloheximide (CHX) chase assay in HEK293T RHNO1-HiBiT cells with or without 18 hours of 2.5 mM HU pre-treatment and monitored the change in HiBiT luminescence. In the absence of treatment, RHNO1 exhibited a significant reduction as soon as 30 min after CHX addition, with a half-life of approximately 1 hour. In contrast, following HU treatment, RHNO1 protein half-life was significantly extended to greater than 6 hours (**Fig. 3D**). Furthermore, following HU washout in the presence of CHX, the loss of DNA replication stress results in the dephosphorylation of RHNO1, shifting the protein back to the unstable unphosphorylated form, which is subsequently degraded. In contrast, continuous HU treatment during CHX chase maintains RHNO1 in its stabilized, phosphorylated form. (**Fig. 3E**). We next investigated if RHNO1 turnover is regulated by the ubiquitin-proteasome system (UPS), by treatment of cells with the proteasome inhibitors MG132 or bortezomib. Both MG132 and bortezomib treatment alone was sufficient to induce RHNO1 protein levels to the levels observed under replication stress, and HU treatment together with either MG132 or bortezomib led to no further increase in RHNO1 levels at 6 hours, while 24 hours post-treatment, combination of HU and either MG132 or Bortezomib further increased RHNO1 protein levels (**Fig. 3F** and **G**). These data support that proteolytic turnover is an important mechanism of RHNO1 regulation, which is shifted under conditions of replication stress.

A previous study reported that CDK1 and PLK phosphorylation of RHNO1 leads to its increased expression during M phase of the cell cycle, whereas only low level expression of RHNO1 was observed in G1/S^21^. As this appears inconsistent with the function of RHNO1 in the ATR/CHK1 replication stress response in S phase, we reexamined endogenous RHNO1 expression throughout the cell cycle using HEK293T RHNO1-HiBiT knock in cells. We performed cell cycle synchronization using a thymidine block (G1/S), CDK1 inhibitor treatment (G2/M), and nocodazole treatment (M), to evaluate RHNO1 expression during specific cell cycle phases (**Supplementary Fig. 5B**). Notably, in this setting, endogenous RHNO1 protein expression was mainly observed during G1/S phase (high cyclin E1/low phosphorylated histone H3 serine 10 (pH3S10)), with very low expression during G2/M and M (low cyclin E1/high pH3S10) (**Supplementary Fig. 5C**). Importantly, although RHNO1 levels were elevated during S phase, it did not exhibit the molecular weight shift observed under replication stress conditions. Collectively, these data suggest that under normal conditions, RHNO1 is mainly expressed in G1/S phase, however, under conditions of replication stress, RHNO1 is phosphorylated, which stabilizes the protein and inhibits its degradation by proteasomes.

### ATR/Chk1-dependent phosphorylation of RHNO1 is required for its stabilization and chromatin recruitment at stalled replication forks

Given that phosphorylation of RHNO1 during replication stress protects it from proteasome degradation (**Fig. 3F**), we next investigated the upstream kinases responsible for RHNO1 phosphorylation. To test this, we treated HEK293T RHNO1-HiBiT cells with a panel of kinase inhibitors (0.2 µM AZD6738 (ATRi); 0.2 µM MK8776 (Chk1i); 9 µM RO-3306 (CDK1i); 50 nM BI2536 (PLK1i); each with or without HU treatment for 24 hours before protein isolation. The single treatment of ATRi or Chk1i did not affect RHNO1 levels or phosphorylation in comparison to HU-treated cells (**Fig. 4A**). Surprisingly, inhibition of CDK1 alone induces the accumulation of unphosphorylated RHNO 1, whereas inhibition of PLK 1 alone induces both accumulation and phosphorylation of RHNO1, comparable to HU treatment (**Fig. 4A**). Importantly Chk1i inhibitor treatment led to a marked reduction of RHNO1 protein expression in combination with HU, while ATRi treatment also led to a lesser reduction (**Fig. 4A**). Importantly, inhibition of CDK 1 or PLK 1 had no effect on RHNO 1 levels or phosphorylation under conditions of replication stress (**Fig. 4A**). These data suggest that ATR/Chk1 activity promotes RHNO1 stabilization during replication stress.

**Figure 4.**
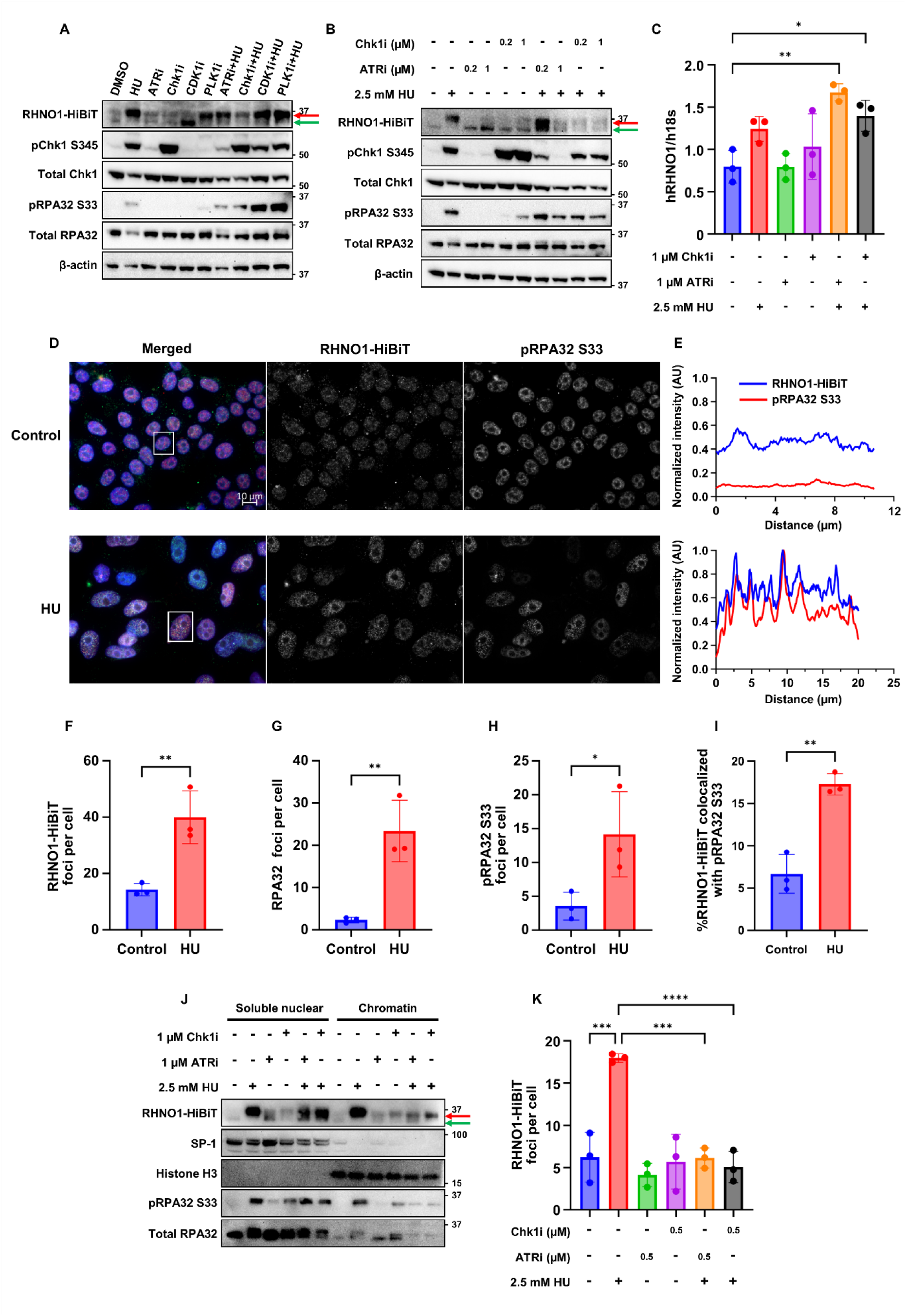
ATR/Chk1 signaling is required for RHNO1 stabilization and localization to stressed replication forks. **A** Western blot analysis of RHNO1-HiBiT in HEK293T RHNO1-HiBiT cells treated with 2.5 mM HU in combination with 0.2 µM AZD6738; ATRi, 0.2 µM MK8776; Chk1i, 9 µM RO-3306; CDK1i, 50 nM BI2536; PLK1i. **B** Effect of low (0.2 µM) and high (1 µM) concentration of ATR inhibitor and Chk1 inhibitor on RHNO1-HiBiT protein expression analyzed by western blotting. **C** RHNO1 mRNA levels in HEK293T RHNO1-HiBiT after 24 hours of 2.5 mM HU incubation with or without ATR and Chk1 inhibitor. **D** Representative immunofluorescence images of RHNO1-HiBiT and pRPA32 S33 foci after 24 hours of 2.5 mM HU treatment. **E** Line scanning analysis of RHNO1-HiBiT and pRPA32 S33 in representative nuclei (white box in **D**). **F – H** Quantification of RHNO1-HiBiT (**F**), RPA32 (**G**), and pRPA32 S33 (**H**) foci in HEK293T RHNO1-HiBiT cell line. **I** Percentage of RHNO1-HiBiT foci colocalized with pRPA32 S33 in HEK293T RHNO1-HiBiT after 24 hours of 2.5 mM HU treatment. **J** Fractionated western blotting of HEK293T RHNO1-HiBiT treated with 24 hours 2.5 mM HU with or without ATR and Chk1 inhibitor. **K** Quantification of RHNO1-HiBiT foci per cell following 24 hours of 2.5 mM HU treatment with or without ATR and Chk1 inhibitor. Data is presented as mean ± SD of at least three independent experiments. Statistical significance was determined using unpaired Student’s t test two-tailed (**F - H**), and one-way ANOVA with Tukey post-hoc test (**C** and **K**) (*p < 0.05, **p < 0.01, ***p < 0.001, ****p < 0.0001). The scale bar represents 10 µm.

We hypothesized that the stronger effect of Chk1i than ATRi on RHNO1 expression during replication stress may be due to the differences in inhibitor potencies. To investigate this, we repeated the experiment using two different concentrations (0.2 µM and 1 µM) of ATR and Chk1 inhibitors, in combination with HU. Consistent with previous findings, inhibition of either ATR or Chk1 alone, regardless of concentration, has no effect on RHNO1 levels or phosphorylation (**Fig. 4B**). However, under replication stress conditions, both 0.2 µM and 1 µM of Chk1 inhibitor reduced RHNO1 expression, while the ATRi at 1 µM concentration also led to significantly decreased RHNO1 protein expression (**Fig. 4B**). Similar findings were also observed when replication stress was induced by etoposide (**Supplementary Fig. 6A**).

The data presented above demonstrates that ATR/Chk1 signaling regulates RHNO1 at the protein level but does not exclude other mechanisms of regulation. To test whether RHNO1 is additionally regulated by ATR/Chk1 signaling at the RNA level, we measured *RHNO1* mRNA in HEK293T RHNO1-HiBiT cells following 24 hours treatment of 2.5 mM HU, with or without ATR/Chk1 inhibitor co-treatment. Although not significant, HU treatment did increase RHNO1 mRNA levels compared to the control, whereas neither single treatment of ATR nor Chk1 inhibitors affected *RHNO1* mRNA levels (**Fig. 4C**). Moreover, the combination of HU with ATR or Chk1 inhibitions did not reduce *RHNO1* mRNA levels, indicating that ATR/Chk1 signaling does not regulate RHNO1 expression at the mRNA level (**Fig. 4C**).

Having established that RHNO1 accumulation and phosphorylation is regulated by ATR/Chk1 during replication stress, we next investigated the recruitment of RHNO1 to genomic sites of replication stress. To visualize the localization of RHNO1 during replication stress, we performed immunofluorescence in HEK293T RHNO1-HiBiT cells after 24 hours treatment of 2.5 mM HU. Under replication stress, we observed significantly increased RHNO1 nuclear foci alongside RPA32 and pRPA32 S33 (**Fig. 4D-H**). Furthermore, under replication stress, we observed significantly increased RHNO1 nuclear foci colocalization with pRPA32 S33 and RPA32 compared to control, indicating the presence of RHNO1 at stalled replication forks (**Fig. 4E** and **I**, **Supplementary Fig. 6B** and **C**).

We next addressed the potential impact of ATR/Chk1 signaling on RHNO1 function during the cellular response to replication stress. Initially, we investigated the effect of ATR/Chk1 inhibition on RHNO1 subcellular localization during replication stress. In the soluble nuclear fraction, RHNO1 protein remained detectable even when ATR/Chk1 signaling was inhibited (**Fig. 4J**). In contrast, in the chromatin-bound fraction, there were marked reductions in RHNO1 levels when ATR or Chk1 inhibitors were co-treated with HU, as compared to HU treatment alone, suggesting that RHNO1 chromatin localization is dependent on ATR/Chk1 activity (**Fig. 4J**). This hypothesis was further supported by immunofluorescence data, which demonstrated significantly reduced RHNO1 nuclear foci in when HU was co-treated with ATR or Chk1 inhibitors (**Fig. 4K** and **Supplementary Fig. 7**). Taken together, these data support that RHNO1 is phosphorylated by ATR/Chk1 signaling during replication stress, and this phosphorylation event promotes RHNO1 stabilization and localization to stressed replication forks.

Finally, we investigated the specific phosphorylation sites that mediate RHNO1 localization under replication stress. We first analyzed RHNO1 amino acid sequence for the minimal Chk1 substrate consensus motif, RXX(S/T)^25^. This analysis identified three putative sites for Chk1 phosphorylation in RHNO1: serine 11, serine 83, and threonine 87. We subsequently performed a rescue assay using wild-type, individual alanine mutated (S11A, S83A, T87A) and all sites mutated (3A) of HA-RHNO1 in a HEK293T RHNO1-HiBiT knock out cell line and quantified the nuclear foci after HU treatment (**Supplementary Fig. 8A-C**). Although not statistically significant, we observed a consistent trend toward reduced RHNO1 and pRPA32 S33 foci in S83A and 3A samples under replication stress conditions (**Supplementary Fig. 8D** and **E**). These findings highlight serine 83 as a candidate residue for ATR/Chk1-mediating RHNO1 localization under replication stress. However, the persistence of RHNO1 foci in the described mutants suggests that other residues and mechanisms also contribute to RHNO1 nuclear foci formation under conditions of replication stress.

## Discussion

DNA replication is a critical cellular process that requires tight regulation in order to suppress genomic instability, especially in the context of elevated replication stress. The ATR/Chk1 signaling pathway is an essential regulator of the replication stress response and both maintains faithful DNA replication and ensures cell survival^5^. Understanding how this pathway (collectively called the DNA checkpoint) is regulated provides insight into genomic maintenance and potential therapeutic vulnerabilities in cancer cells^5,9^. While the mechanism of initial activation of the ATR/Chk1 pathway is well established, what mechanisms sustaining the initial signal is less understood. In this study we provide the first experimental evidence that RHNO1 functions as more than just a scaffold for ATR/Chk1 activation, but also as an integral part of the signaling required to sustain the DNA checkpoint^15,16^.

Our data reveal a distinction between the initiation and the maintenance of ATR/Chk1 signaling upon replication stress. RHNO1 was first identified as a scaffolding protein for ATR/Chk1 initiation; however, during the acute phase of replication stress, we observed normal ATR/Chk1 activation even in the absence of RHNO1. This suggests that the initial activation of ATR/Chk1 may rely solely on the canonical 9-1-1/TopBP1 complex^26^. However, we demonstrate that, in the absence of RHNO1, cancer cells fail to sustain the signal at later time points after replication stress. These data agree with our recent report that mouse cells sustaining *Rhno1* deletion have defective resolution of radiation induced DNA damage specifically at later time points^27^.

Mechanistically, RHNO1 protein is kept at very low levels due to proteolytic degradation under basal conditions but is stabilized and upregulated after replication stress. Structurally, RHNO1 has been reported to bind to Rad1 in the 9-1-1 complex^17^. It is possible that phosphorylation of RHNO1 induces conformational changes that either mask a degron motif of RHNO1 or increase its binding affinity to Rad1, forming a stable 9-1-1/RHNO1 heterotetramer, shielding it from proteasomal degradation. We propose that this instability of RHNO1 serves as a “kill switch” for the ATR/Chk1 signal termination once replication stress is resolved. However, under replication stress, RHNO1 protein is needed to maintain the signal; therefore, the degradation is halted. We found that

ATR/Chk1 signaling is required for stabilization of RHNO1 during replication stress, via phosphorylation. Furthermore, our data suggests that this phosphorylation facilitates RHNO1 accumulation at stalled replication forks. Inhibition of ATR or Chk1 signaling during replication stress was found to prevent stabilization of RHNO1 protein and to abolish both its chromatin localization and its presence in nuclear foci.

Collectively, our findings suggest that RHNO1 is a downstream effector of ATR/Chk1, not only a scaffold for signal activation^15,16^. Notably, this temporal distinction of RHNO1 function is in agreement with a recent computational modeling study, which demonstrated that the initial ATR signal activation is driven by Rad17-mediated loading of the 9-1-1 complex to the damaged DNA and the interaction of Rad9 tail with TopBP1, both independent of RHNO1^26,28^. The modeling study proposed that after the initial ATR signal activation, RHNO1 might replace Rad17 binding to the 9-1-1 complex and facilitate the bridging of multiple 9-1-1 complexes on damaged DNA, sustaining the initial ATR signal activation^28^. This bridging function of RHNO1 is dependent on second KYxxL+ motif with Arg84 and Lys90 as a critical residue^28^. Our data has identified Ser83 as a possible Chk1 phosphorylation site required for RHNO1 localization at the DNA and this residue is adjacent to Arg84. Therefore, it is possible that phosphorylation of RHNO1 by Chk1 at Ser83 during replication stress response may help to facilitate the bridging function of RHNO1, preventing premature ATR/Chk1 signal collapse.

Based on these collective findings, we propose a novel model whereby dynamic regulation of RHNO1 creates a positive feedback loop where the initial ATR/Chk1 activated under replication stress stabilizes RHNO1, which is then required to sustain the checkpoint signaling at stressed replication forks (**Fig. 5**). This positive feedback loop may help ensure that the downstream signaling of ATR/Chk1 is maintained until replication stress is resolved, preventing premature mitotic entry and thus genomic instability.

**Figure 5.**
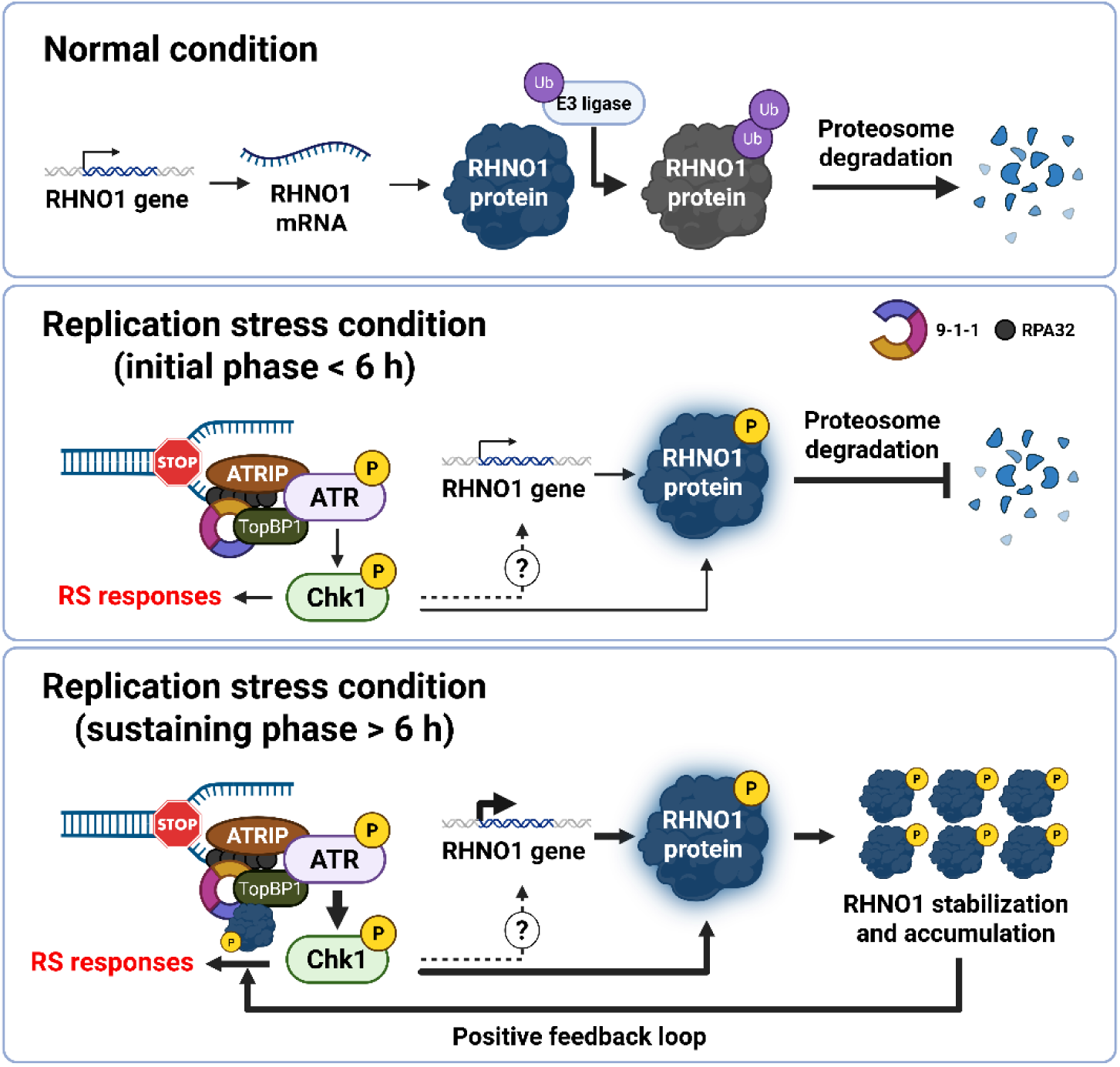
Proposed model of the RHNO1 and ATR/Chk1 positive feedback loop. Schematic illustrating RHNO1 regulation under normal and replication stress conditions. Under normal conditions, RHNO1 is unstable and rapidly degraded by the proteasome. Upon replication stress (initial phase), ATR/Chk1 signaling is activated independently of RHNO1, however, this signal phosphorylates RHNO1, preventing its proteasomal degradation. In the sustaining phase, the RHNO1 started to accumulate and localized at the stalled replication forks to maintain and sustain ATR/Chk1 signaling, creating a positive feedback loop essential for genomic stability and cell survival under replication stress.

The regulation of RHNO1 expression reported here appears to be analogous to other critical cell cycle-related proteins such as CDKs, cyclins, or Cdc25A, which are tightly regulated by phosphorylation and proteasome-mediated degradation^24^. Consistent with this temporal control, we previously demonstrated that the RHNO1 gene shares a bidirectional promoter with the oncogene FOXM1, a critical transcription factor involved in cell cycle control^18^. The bidirectional gene arrangement and tight co-expression of FOXM1 and RHNO1 suggests intrinsic coordination with active cell division, ensuring that RHNO1 is expressed during the stages of the cell cycle where replication stress arises, particularly during S phase. Furthermore, a recent study identified D-box and KEN-box motifs, an anaphase-promoting complex (APC/C) recognition sequence in RHNO1^21^. These motifs were shown to regulate RHNO1 expression during M phase, where it facilitated TMEJ^21^. The role for RHNO1 in mitotic DNA repair is also supported by our data in *Rhno1* knockout B cells^27^. Additionally, we also observed that RHNO1 knockdown in ovarian cancer cells leads to TMEJ impairment.

In contrast to the presumed mitotic function of RHNO1 in TMEJ^21^, our data using the HEK293T RHNO1-HiBiT knock in cell model demonstrated that RHNO1 expression was mainly observed during G1/S phase, where APC/C is less active^29^. While the inactivation of APC/C may contribute to RHNO1 accumulation, our data strongly suggest that basal ATR/Chk1 signaling in response to intrinsic replication stress also plays a role^30^. This notion is also supported by a recent study suggesting that the 9-1-1 complex, which physically interacts with RHNO1, mediates replication fork protection by blocking EXO1 nucleolytic of single-stranded DNA and promoting DNA polymerase ζ-dependent gap filling^31^. Therefore, it is possible that intrinsic ATR/Chk1 signaling may be sufficient to stabilize RHNO1 to mediate both the replication stress response, including fork protection. Consequently, the micronuclei formation observed in RHNO1-depleted OVCAR8 cells we report here may result both from failure to resolve intrinsic replication stress during S phase as well as TMEJ in M phase^21,30^.

In addition to this mechanistic information, we verified the importance of RHNO1 in cancer biology using ovarian cancer cell lines and xenografts. Tumors sustaining RHNO1 depletion showed reduced growth and ascites, and increased mouse survival. Notably, ovarian cancer has high intrinsic DNA replication stress due to defective DNA damage repair machinery and oncogene overexpression^4,18^. We hypothesize that disrupting the RHNO1-ATR/Chk1 positive feedback loop may increase genomic instability in cancers with heightened basal replication stress, causing cell death due to increased DNA damage. Consistently, other recent studies also have pinpointed a role for RHNO1 in cancer biology and prognosis, including in liver and lung cancer^32–34^. Furthermore, beyond its role in the replication stress response, RHNO1 may contribute to other cell survival pathways such as PI3K/Akt and NF-kB, and thus targeting RHNO1 may have other important anti-tumor mechanisms beyond ATR/Chk1 signaling^18,32–34^. It is additionally worth noting that Rhno1 knockout mice are viable, suggesting limited functions for *Rhno1* in normal tissues, further suggesting the therapeutic value of RHNO1 as a cancer therapeutic target^27^.

Finally, we note that recent high throughput screens have identified RHNO1 as a potential synthetic lethal candidate in BRCA2-deficient cells^35,36^. Our findings are in agreement with this observation; in the absence of BRCA2-mediated homologous recombination, cancer cells rely heavily on RHNO1 and ATR/Chk1 signaling to counteract excessive replication stress. In summary, RHNO1 is an interesting potential target for therapeutic intervention in cancers with BRCA2 deficiency and those that exhibit elevated replication stress.

## Methods

### Cell cultures

OVCAR8 cell line was obtained from the National Cancer Institute Division of Cancer Treatment and Diagnosis Cell line repository, and HEK293T cell line was obtained from American Type Culture Collection (ATCC). All cell lines were maintained in DMEM (Corning) supplemented with 5% FBS, 100 U/ml penicillin, and streptomycin, and 2.5 µg/ml Plasmocin^®^ Prophylactic (InvivoGen). Cell lines were regularly screened for mycoplasma contamination using a nested PCR method.

### Lentiviral packaging and transduction

The lentiviral packaging protocol was performed as previously described^18^. The lentiviral titer was determined using qPCR Lentivirus Titer Kit (Applied Biological Materials). OVCAR8 stably expressing non-silencing (NS) or RHNO1 specific shRNA (RHNO1-130 and -133) under a pGIPZ backbone were generated by lentiviral transduction. The transduced cells from each constructed were selected by puromycin and isolated for monoclonal line. The presence of RHNO1 knockdown was confirmed by qPCR. OVCAR8 GIPZ cell lines with stably expressed firefly luciferase were generated by lentiviral transduction, selected with G418, and isolated for monoclonal line. The luciferase activity was measured by GLOMAX luminometer (Promega).

### CRISPR-Cas9 gene editing

Guide RNA targeting C-terminal sequence of RHNO1 was cloned into pLentiCRISPRv2 blasticidin (Addgene #83480) according to Zhang’s lab protocol. The knock-in homology template plasmid harbored glycine-serine (GS) linker, and HiBiT tag sequences was synthesized and designed by Alt-R™ HDR Design Tool (IDT). The plasmids were transfected into HEK293T cell line followed by blasticidin selection, and monoclonal isolation. The clone with successful knock-in was confirmed by genomic DNA PCR using primers flanking the homology arms. To generate HEK293T RHNO1-HiBiT knock-out cells, gRNA was designed using CHOPCHOP^37^ and cloned into pLentiCRISPRv2 blasticidin. The plasmid was transfected into HEK293T RHNO1-HiBiT cells, selected with blasticidin, and isolated for monoclonal line. The successful RHNO1 knock-out was confirmed by PCR and western blotting. All gRNA and primer sequences used are listed in **Supplementary Table 1**.

### Clonogenic growth assays

Cells were plated at a density of 500 cells in triplicate wells of 6-well plate and cultured for 10-14 days. Following incubation, cells were fixed in ice-cold methanol and stained with 0.5% crystal violet in PBS/methanol, rinsed with water, and air-dried overnight. The plates were photographed and colonies containing more than 50 cells were counted using Fiji ImageJ software (NIH). For soft-agar clonogenic assay, 2,500 cells suspended in 0.4% agarose were plated on top of solidified 0.8% agarose in 6-well plate. After the gel solidified, 1 ml of culture media was added on top of the agarose gel and cultured for 14 days. After incubation, soft agar was stained with 0.01% crystal violet in 10% ethanol. Rinsed with water and air-dried overnight. The plates were photographed and colonies containing more than 50 cells were counted using countPHICS software^38^.

### Cell viability assays

For cell growth monitoring experiment, 500 cells were seeded into triplicate wells of a 96-well plate. The cell viability was monitored by Alamar Blue reagent (BioRad) every 24 hours for 7 days. For the HU sensitivity assay, 2,000 cells were seeded into triplicate wells of 96-well plate. Cells were treated with HU for 72 hours before replacing with fresh media for another 48 hours before cell viability measurement using Alamar Blue. The excitation and emissions wavelength for Alamar Blue are 560 and 590 nm, respectively.

### Reverse transcription quantitative PCR (RT-qPCR)

Total RNA was isolated using TRIzol (Invitrogen) and Direct-zol RNA Purification Kit (Zymo Research) following the manufacturer’s instructions. A total of 0.5 – 1 µg of RNA was converted to cDNA using the iScript™ cDNA Synthesis Kit (BioRad). cDNA was diluted 1:5 and used as template for qPCR reaction using iScript™ Reverse Transcription Supermix for RT-qPCR (BioRad) following the manufacturer’s instructions. The qPCR was performed in triplicates in the CFX Connect Real-Time System (BioRad). Relative gene expression was normalized using *h18s* as a housekeeping gene, and all calculations were performed using the Δ ΔCt method. A list of the primers sequences is provided in **Supplementary Table 1**.

### Immunofluorescence

1 x 10^5^ Cells were plated on a coverslip and incubated with treatments as indicated. All immunofluorescence was performed with pre-extraction buffer (20 mM HEPES pH 7.5, 50 mM NaCl, 3 mM MgCl_2_, 300 mM sucrose, 0.5% Triton-X 100) incubation for 1 min on ice before cells fixation except in the micronuclei quantification experiment. The cells were fixed with 4% formaldehyde for 10 minutes followed by 0.5% Triton-X 100 permeabilization for 10 minutes. Cells were then blocked for 30 minutes using blocking buffer (5% BSA dissolved in 0.1% Triton-X 100 in PBS). Cells were incubated with primary antibodies (1:400 dilution) in blocking buffer for 1 hour followed by secondary antibodies (1:1000) incubation in blocking buffer for 30 minutes. Cells were mounted with VECTASHIELD® Vibrance™ Antifade Mounting Medium with DAPI (Vector Laboratories). The images were captured using ZEISS Axio Observer Z1/7 confocal microscope at 63x magnification. Foci detection and colocalization analysis were performed using Fiji ImageJ (NIH). Binary masks were generated for each channel, and foci were identified using particle analysis and added to the ROI Manager. Colocalization was quantified in a nucleus-restricted manner by assessing the overlap between RHNO1 and pRPA32 foci using a custom ImageJ macro. For HA-RHNO1 rescued experiment, HEK293T RHNO1-HiBiT KO cells were plated into 6-wells plate and transfected with 150 ng of pCMV6-HA-RHNO1 (WT or mutant) using lipofectamine 3000. 24 hours after transfection, cells were sub-cultured to coverslip and treated with 2.5 mM HU for 24 hours before immunofluorescence staining as described. The list of antibodies used is provided in **Supplementary Table 1**.

### Chromosomal assay to measure TMEJ

Double strand breaks (DSBs) were generated by Cas9 in the human *LBR* gene locus by electroporation of RNP complex of 36 pmol Alt-R crRNA annealed to Alt-R tracrRNA (IDT) and 30 pmol Cas9, into 600,000 cells in a 25 μl volume using the Amaxa (Lonza) 4D nucleofector X unit with the SK-OV-3 cell line program (pulse code FE 132). Cells were then incubated for 48 hours before DNA was extracted using the QIAamp DNA mini-kit (Qiagen). Recovered DNA was then analyzed using qPCR primers and probes specific to products dependent TMEJ, or to the break site with no insertions and deletions. We additionally corrected for differences in sample recovery using qPCR amplicon located within the same locus but 10 kilobase pairs distal to the break site. The spacer sequence used for the Alt-R crRNA and all primer and probe sequences are in **Supplementary Table 1**. ART558 was a gift from Artios UK.

### Mouse xenograft experiment

Animal experiments were performed in accordance with protocols approved by the Institutional Animal Care and Use Committee (IACUC) of the University of Nebraska Medical Center (Protocol 24-061-11-FC). Mouse xenograft experiments were conducted using 7-week-old female NU/J homozygous athymic mice (strain 002019) purchased from The Jackson Laboratory. A total of 15 mice were randomized into three groups (n = 5 mice per group). Mice were intraperitoneally injected with 5 × 10^6^ OVCAR8 GIPZ NS, RH130, or RH133 cells stably expressing luciferase. Tumor growth was monitored twice weekly via bioluminescence imaging. Briefly, mice were intraperitoneally injected with 150 mg/kg D-luciferin substrate (XenoLight) and the bioluminescence signal was captured using the PerkinElmer IVIS Spectrum In Vivo Imaging System (PerkinElmer). At the experimental endpoint, mice were euthanized via CO_2_ asphyxiation followed by cervical dislocation.

### Western blotting

The whole cell lysates were isolated using M-PER^TM^ protein lysis buffer (Thermo Fisher) supplemented with protease and phosphatase inhibitors. For soluble and chromatin-bounded protein isolation, Subcellular Protein Fractionation Kit for Cultured Cells (Thermo Fisher) were used according to the manufacturer’s protocol. To prepare λ phosphatase-treated cell lysates, the whole cell lysates were isolated using M-PER^TM^ lysis buffer with protease inhibitor followed by 400 U λ phosphatase (SantaCruz) incubation according to the manufacturer’s instruction. The total protein concentration was determined using BCA assay (Thermo Fisher). Total protein of 20-40 µg was resolved on 4-12% NuPAGE^TM^ Bis-Tris gel (Thermo Fisher) and transferred to 0.45 µm PVDF membrane (Sigma) using wet transfer protocol. The membranes were blocked with 5% skim-milk and incubated in the primary antibodies overnight followed by HRP-conjugated secondary antibodies incubation for additional hour. The chemiluminescent signal was detected using SuperSignal^TM^ West Pico PLUS substrate (Thermo Fisher) in ChemiDoc Imaging System (BioRad). The protein quantification was calculated using Fiji ImageJ software (NIH). All antibodies used in this study are listed in **Supplementary Table 1**.

### HiBiT lytic luminescence assay

RHNO1-HiBiT luminescence signal was measured using Nano-Glo^®^ HiBiT Lytic Detection System (Promega) according to the manufacturer’s instruction. Briefly, 1-2 x 10^5^ cells were collected after the treatment and incubated in lysis buffer supplemented with 1:100 recombinant LgBiT and 1:50 substrate for 10 minutes followed by luminescence detection using GloMax^®^ 20/20 Luminometer (Promega). The luminescence signal is presented as relative light unit (RLU).

### Cell synchronization experiments

1.5 x 10^6^ HEK293T RHNO1-HiBiT cells were synchronized in G1/S phase by incubating in 2 mM thymidine for 24 hours followed by 16 hours incubation of 9 µM RO-3306 to synchronize the cells to G2/M phase, cells were then released into 100 ng/ml nocodazole for 2 hours to obtain synchronized M phase cells. Cell lysates were collected at different cell cycle phases and investigated for the protein expression using western blotting.

### Cycloheximide-chase assay

1 x 10^6^ HEK293T RHNO1-HiBiT cells were pre-treated with either PBS or 2.5 mM HU for 18 hours before the addition of 10 µg/ml cycloheximide. Cells were harvested at 0, 30 min, 1, 2, 4, and 6 hours after cycloheximide addition to measure RHNO1-HiBiT protein levels using HiBiT lytic luminescence assay or western blotting. The % RHNO1 remaining was calculated as (luminescence signal at each timepoint/luminescence signal at 0 hour) x 100.

### Drug treatments

The list of drugs and inhibitors used in this study is given in the **Supplementary Table 1**. Specific drug concentrations and incubation time used in experiments are described in the figures and legends.

### Statistical analysis

Unpaired Student’s t test was used to compare the difference between two groups. One-way ANOVA with Tukey post-hoc was used to compare the difference between more than two groups. Two-way ANOVA was used to compare the difference in dataset with two variables. Fisher’s exact test was used to assess the difference in ascitic formation incident between groups. The difference in *in vivo* tumor growth rate was determined by Two-way ANOVA mixed effect model. Overall survival of xenograft mouse was calculated by the Kaplan-Meier method, and the significance was determined using the Log-rank test. All data are presented as mean ± standard deviation (SD) from at least three independent experiments unless otherwise stated. All statistical analyses were performed using GraphPad Prism (GraphPad Software, Inc). p value of less than 0.05 was considered statistically significant. Significance levels are indicated in the figures as follows: *p < 0.05, **p < 0.01, ***p < 0.001, and ****p < 0.0001.

## Acknowledgements

This work was supported by NIH R21CA273399 (ARK), US Department of Defense HT9425-23-1-0238 (ARK), NIH P30CA036727 (ARK, GG), NIH R01GM141232 (GG), NIH R01CA263504 (GG, ARK), NCI P01CA247773 (DAR), and NCI P01CA250957 (SFB).

---

**Supplementary figure 1.**
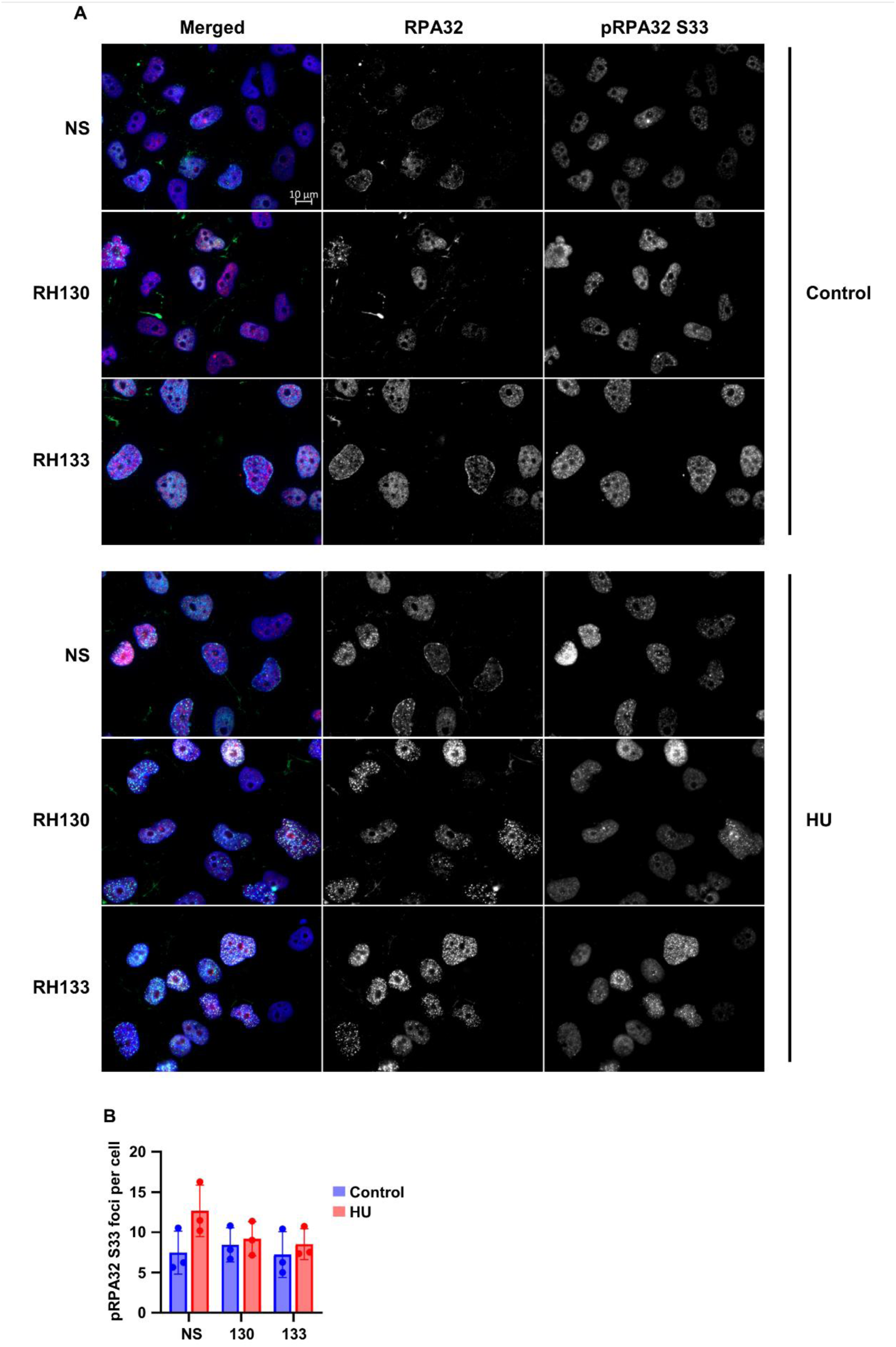
RPA32 and pRPA32 S33 foci in RHNO1-depleted OVCAR8 cell lines. **A** Representative immunofluorescence images of DAPI-stained nuclei showing RPA32 and pRPA32 S33 foci in OVCAR8 NS, RH130, and RH133 cells under control and 2.5 mM HU treatment for 5 h followed by 24 h recovery. **B** Quantification of pRPA32 S33 foci per cell. Data is presented as mean ± SD of at least three independent experiments. The scale bar represents 10 µm.

**Supplementary figure 2.**
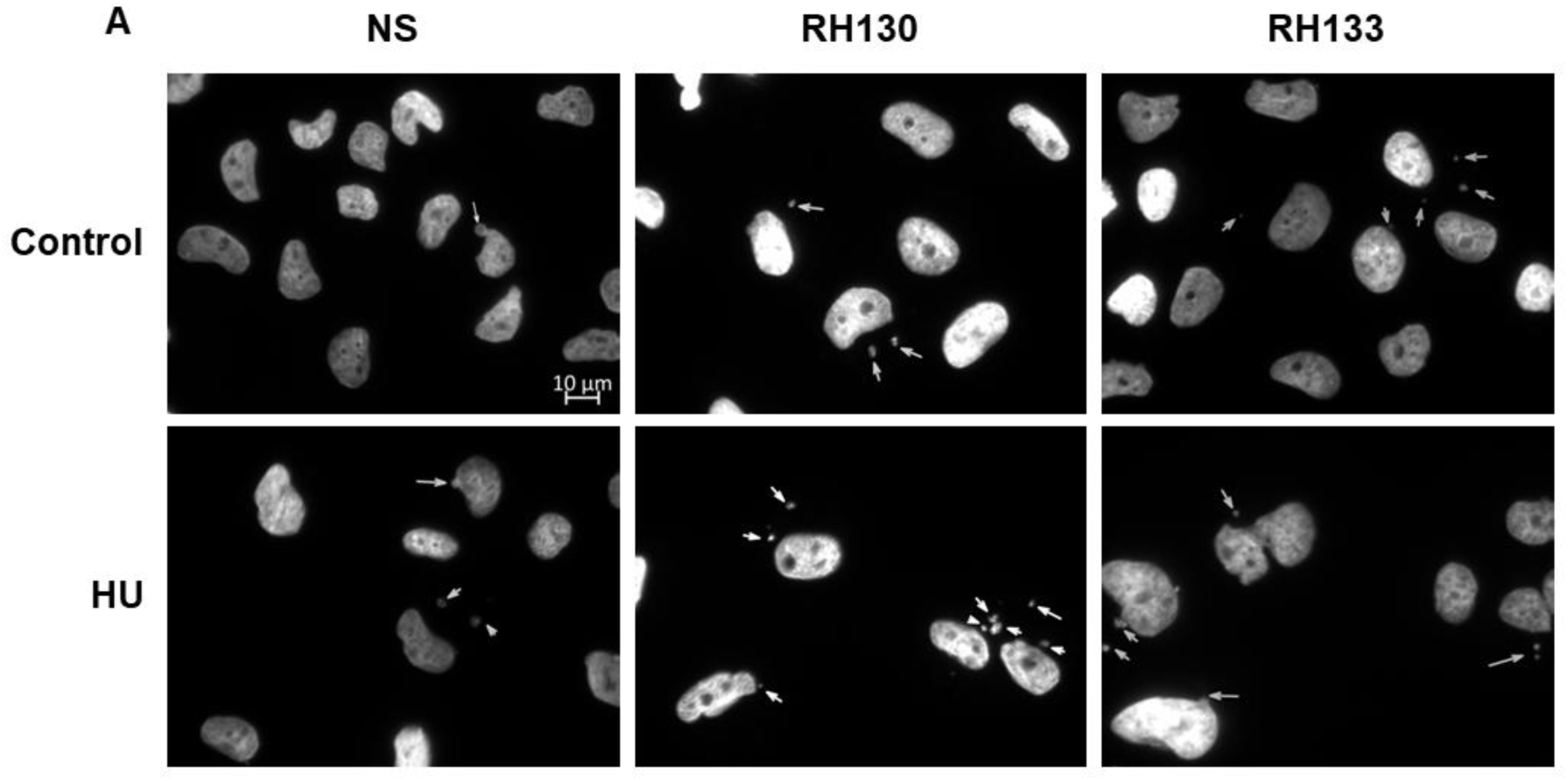
Micronuclei formation in RHNO1-depleted OVCAR8 cell lines. **A** Representative immunofluorescence images of DAPI-stained nuclei showing micronuclei (indicated by arrows) in OVCAR8 NS, RH130, and RH133 cells under control and 2.5 mM HU treatment for 5 h followed by 48 h recovery. The scale bar represents 10 µm.

**Supplementary figure 3.**
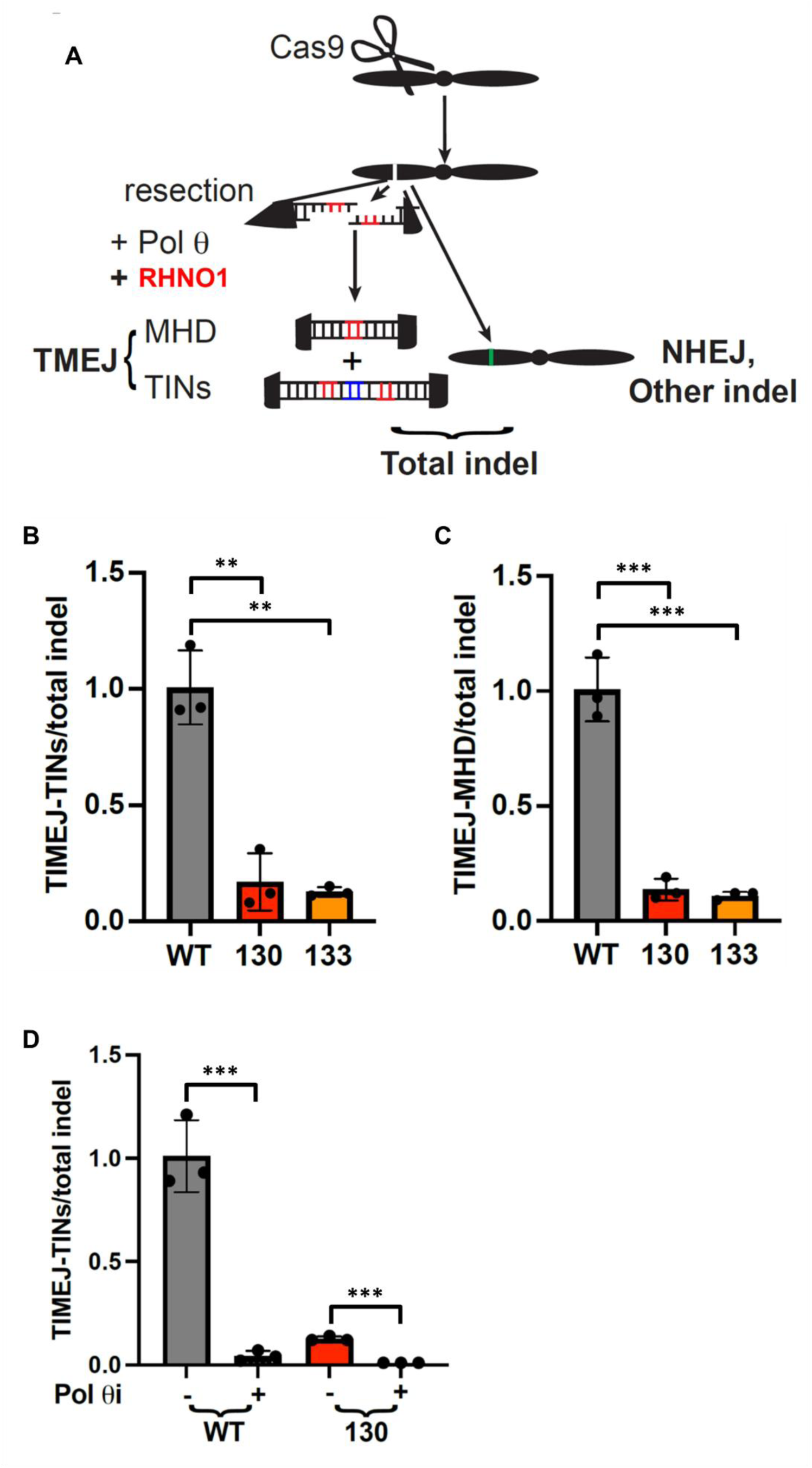
TMEJ activity in RHNO1-depleted OVCAR8 cell lines. **A** Schematic of chromosomal repair assay. Microhomology deletion (MHD) and templated insertions (TINs), as well as total insertions and deletions (indel) were measured by qPCR of DNA recovered 48 hours later after DNA double strand break induction using Cas9 at LBR gene locus. **B** and **C** The frequencies of TMEJ pathway product MHD (**B**) and TINs (**C**) normalized by total indel in OVCAR8 cell lines. **D** Quantification of TINs product in presence of DNA polymerase θ specific inhibitor, ART558. Data is presented as mean ± SD of three independent experiments.

**Supplementary figure 4.**
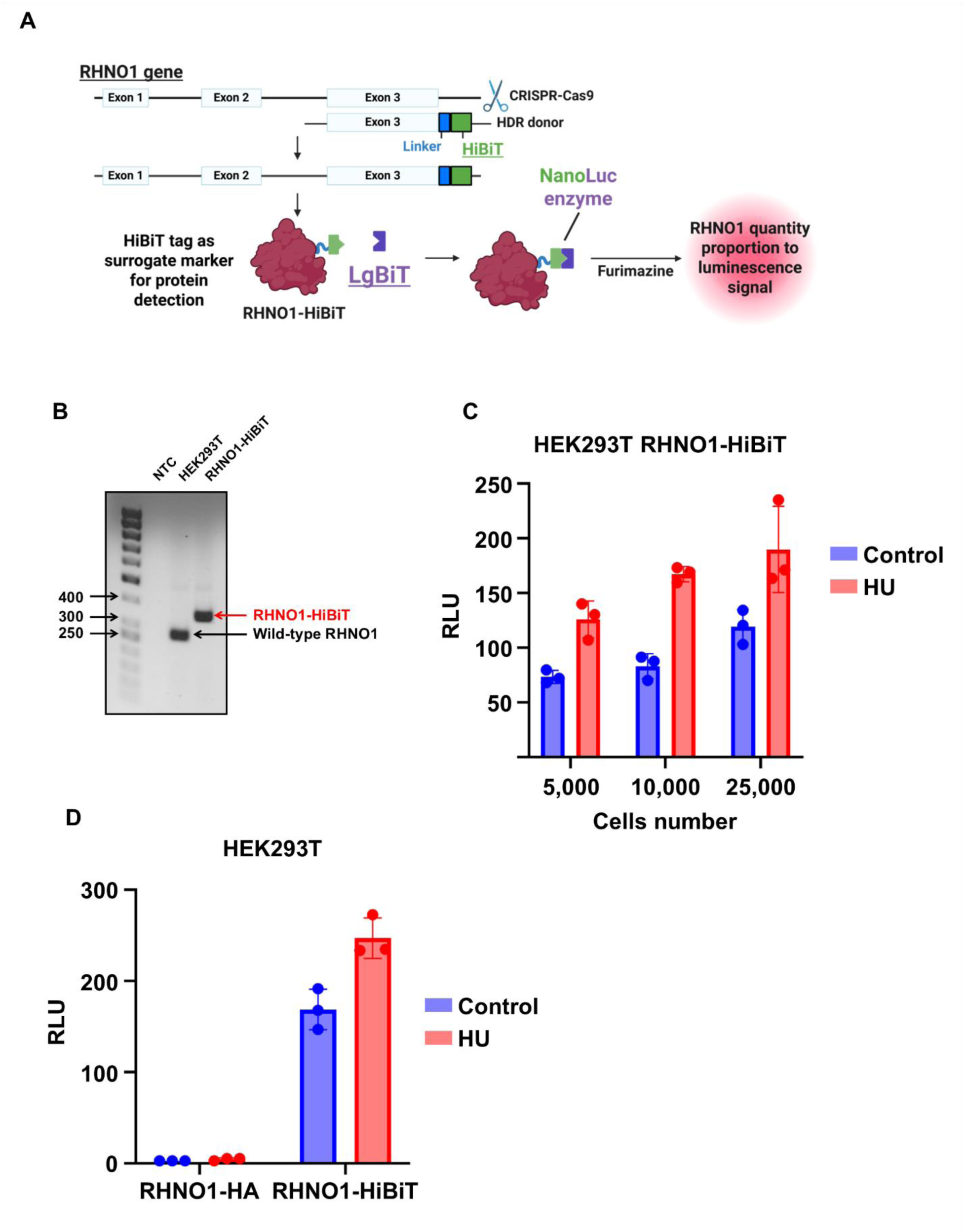
Generation and validation of HEK293T RHNO1-HiBiT cell line. **A** Schematic of CRISPR-Cas9 knock-in strategy for HiBiT-tagged endogenous C-terminal RHNO1. **B** Genomic PCR for the presence of HiBiT tag in RHNO1 gene. **C** HiBiT luminescence signal in HEK293T RHNO1-HiBiT in different concentration of cells. **D** Comparison of HiBiT luminescence signal between RHNO1-HA and RHNO1-HiBiT tagged cells under replication stress conditions. Data from a single experiment with three technical replicates.

**Supplementary figure 5.**
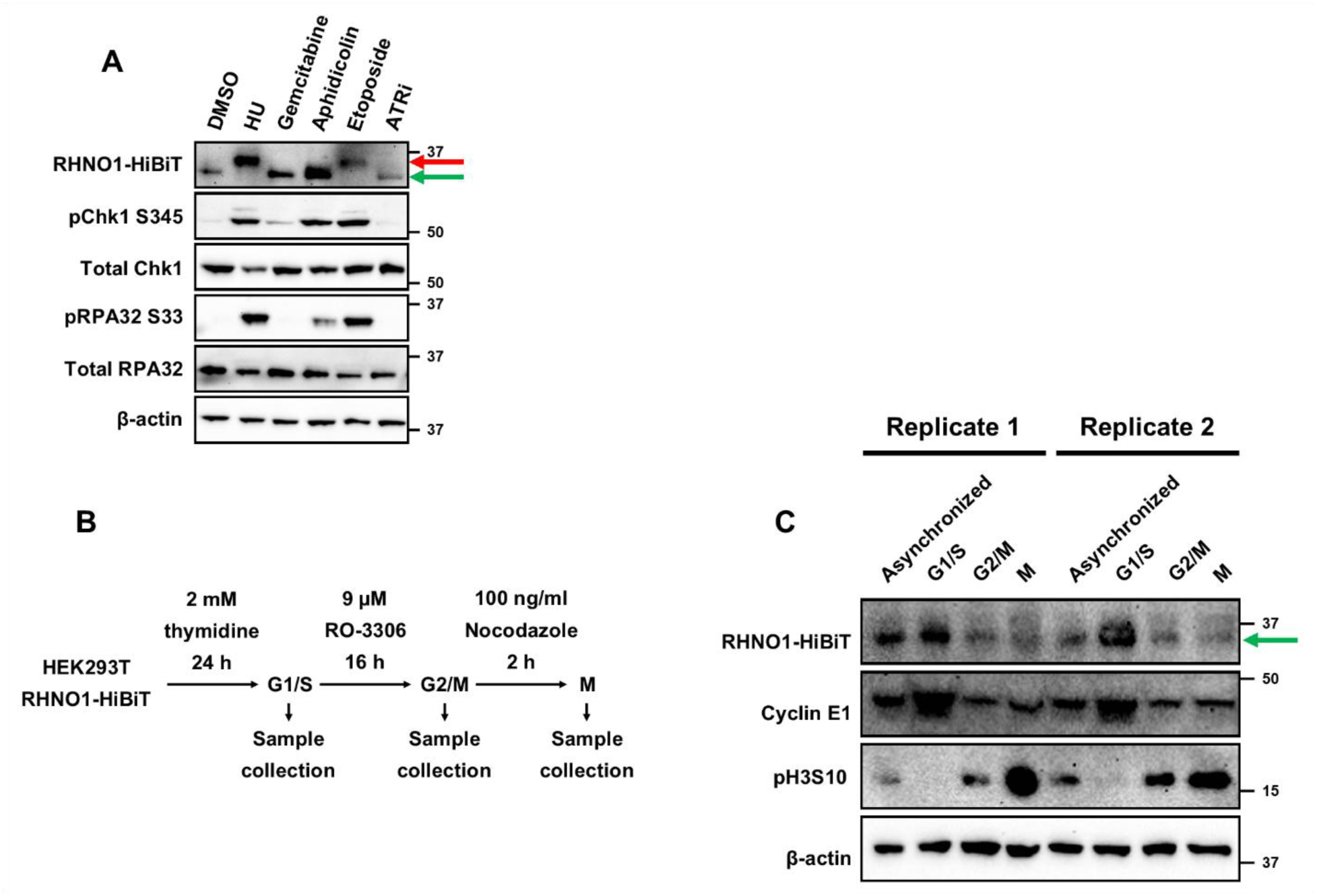
RHNO1 is phosphorylated in response to DNA replication stress inducers and mainly expressed during G1/S phase of cell cycle. **A** Western blotting of RHNO1-HiBiT protein in response to a panel of DNA damaging agents for 24 hours. **B** Schematic of cell cycle synchronization experiment using thymidine, RO-3306, and nocodazole. **C** Western blot analysis of RHNO1-HiBiT protein in different cell cycle phases.

**Supplementary figure 6.**
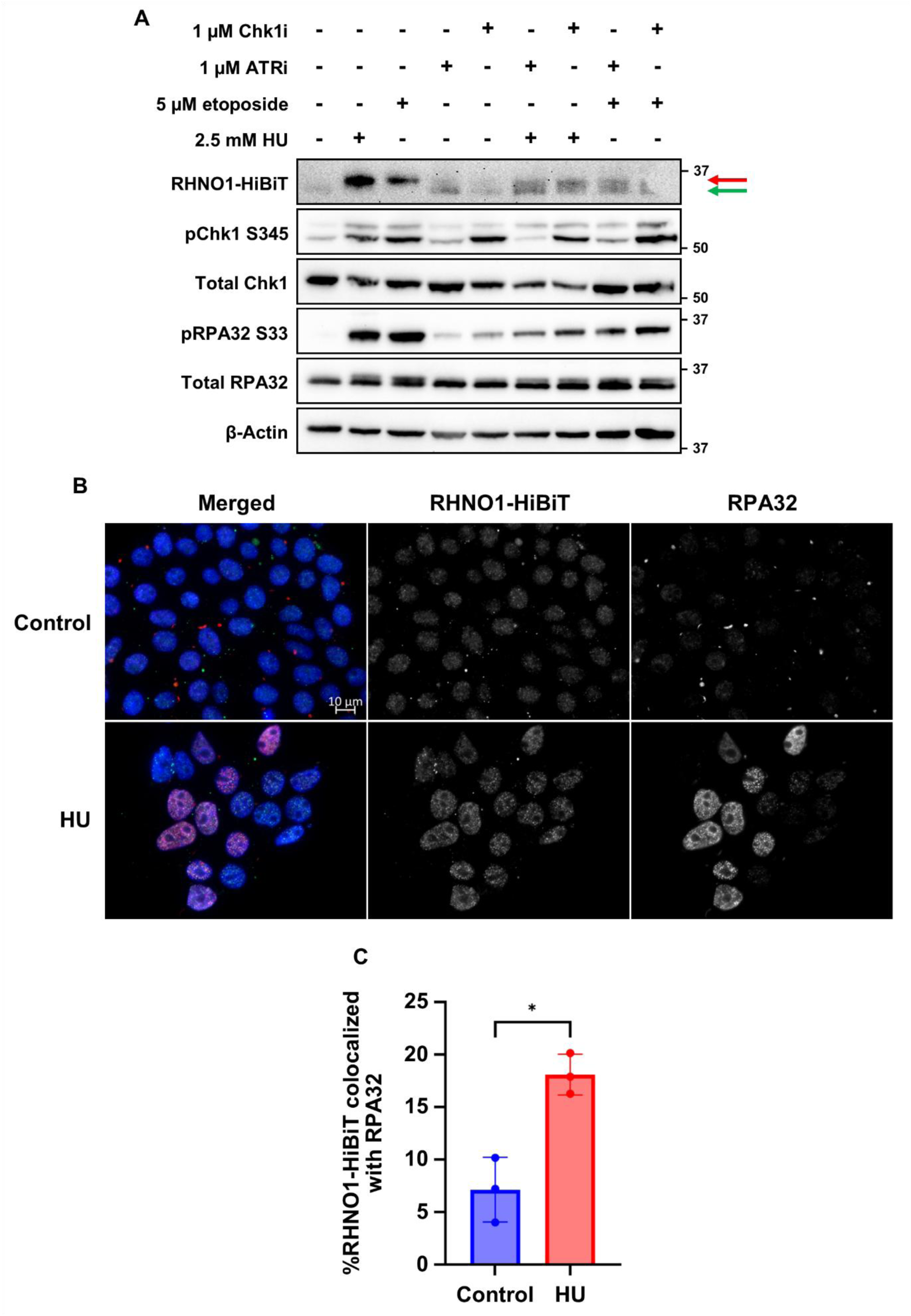
Phosphorylation of RHNO1 under replication stress condition is dependent on ATR/Chk1 kinase activities and RHNO1 form foci and colocalizes at stressed replication forks during stress condition. **A** Western blot analysis of RHNO1-HiBiT in HEK293T RHNO1-HiBiT cells treated with 2.5 mM HU or 5 µM etoposide with or without ATR or Chk1 inhibitors. Phosphorylated and unphosphorylated RHNO1 are noted by red and green arrows, respectively. **B** Representative immunofluorescence images of RHNO1-HiBiT and RPA32 foci in HEK293T RHNO1-HiBiT treated with 2.5 mM HU for 24 h. **C** Colocalization analysis of RHNO1-HiBiT and RPA32 foci in HEK293T RHNO1-HiBiT cell line. Data is presented as mean ± SD of at least three independent experiments. Statistical significance was determined using unpaired Student’s t test two-tailed (*p < 0.05, **p < 0.01). The scale bar represents 10 µm.

**Supplementary figure 7.**
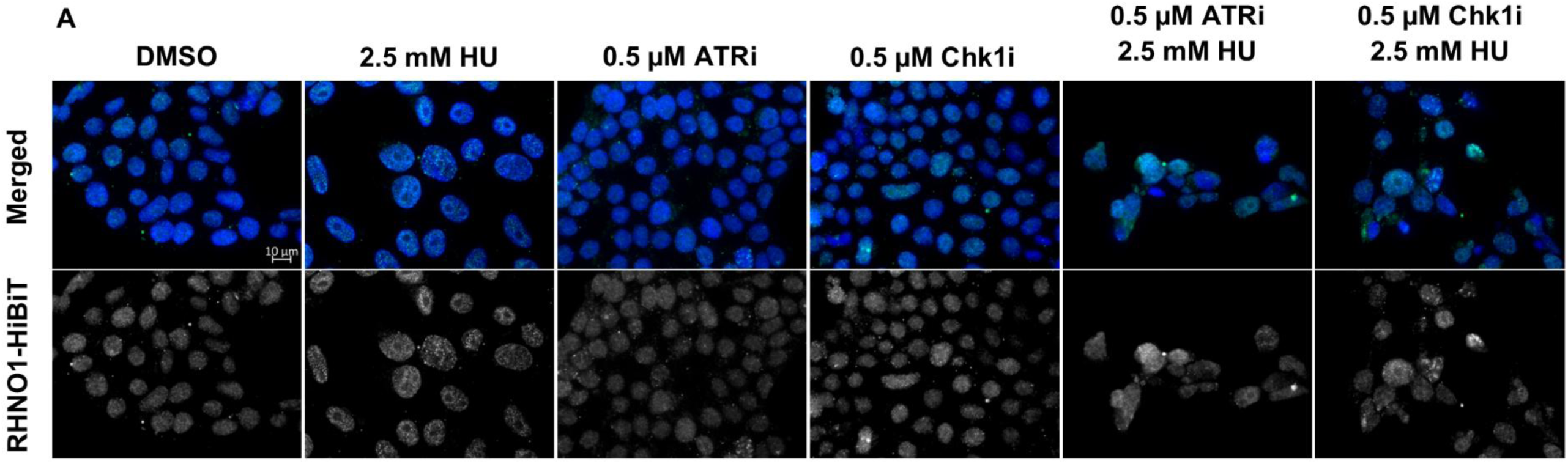
Inhibition of ATR/Chk1 disrupts RHNO1 foci formation under replication stress. **A** Representative immunofluorescence images of HEK293T RHNO1-HiBiT treated with HU alone or in combination with ATR or Chk1 inhibitors. The scale bar represents 10 µm.

**Supplementary figure 8.**
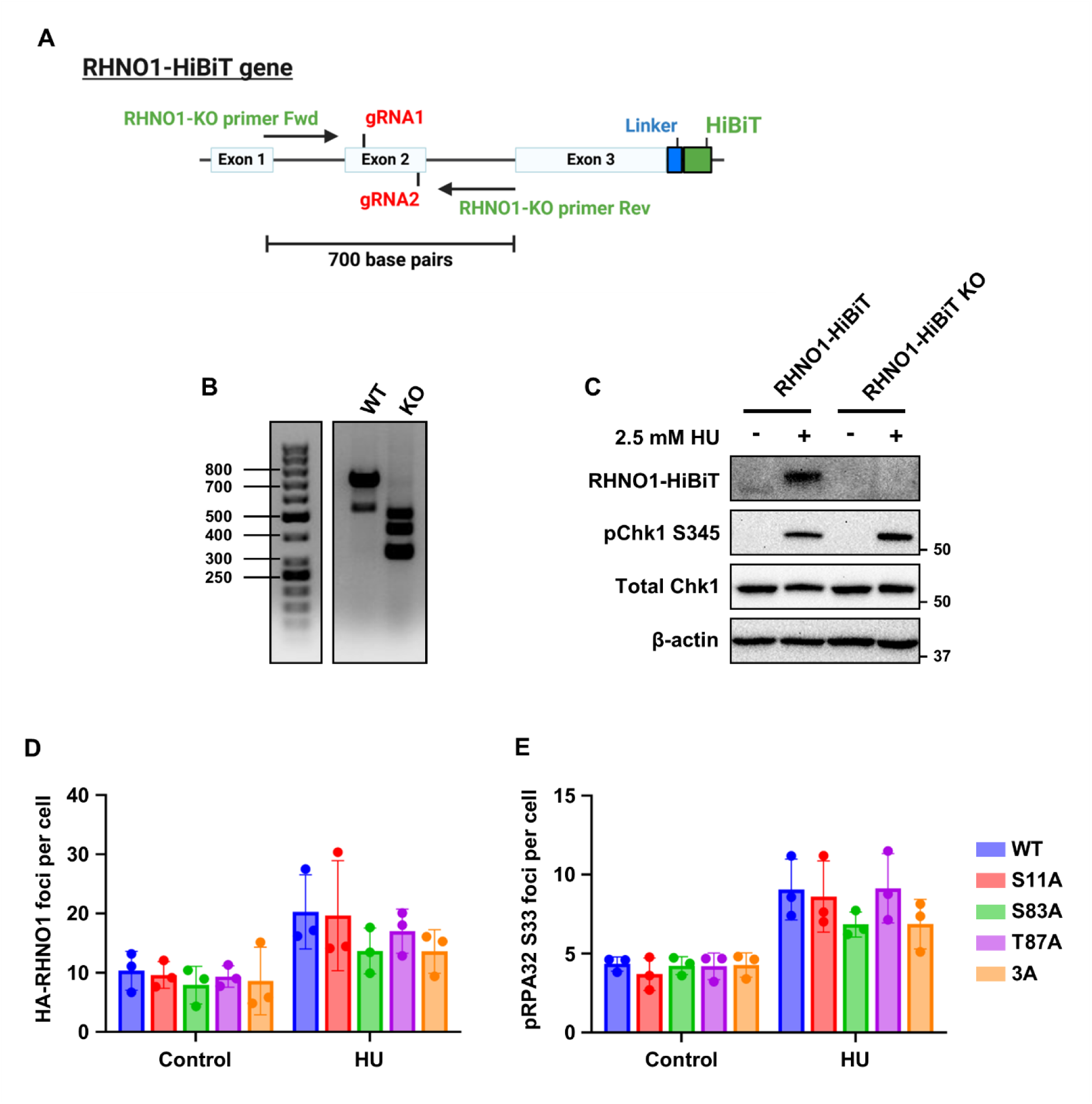
HA-RHNO1 re-expression in HEK293T RHNO1-HiBiT knock out cells. **A** Schematic of CRISPR-Cas9 RHNO1 knock out in HEK293T RHNO1-HiBiT cells. **B** Genomic DNA PCR analysis of parental and knock out cells. **C** Western blotting of RHNO1-HiBiT protein in parental and knock out cells. **D** and **E** quantification of HA-RHNO1 (**D**) and pRPA32 S33 foci (**E**) after 24 hours treatment of 2.5 mM HU. Data is presented as mean ± SD of at least three independent experiments.

